# ChIA-PIPE: A fully automated pipeline for ChIA-PET data analysis and visualization

**DOI:** 10.1101/506683

**Authors:** Daniel Capurso, Jiahui Wang, Simon Zhongyuan Tian, Liuyang Cai, Sandeep Namburi, Byoungkoo Lee, Harianto Tjong, Zhonghui Tang, Ping Wang, Chia-Lin Wei, Yijun Ruan, Sheng Li

## Abstract

ChIA-PET enables the genome-wide discovery of chromatin interactions involving specific protein factors, with base-pair resolution. Interpreting ChIA-PET data depends on having a robust analytic pipeline. Here, we introduce ChIA-PIPE, a fully automated pipeline for ChIA-PET data processing, quality assessment, analysis, and visualization. ChIA-PIPE performs linker filtering, read mapping, peak calling, loop calling, chromatin-contact-domain calling, and can resolve allele-specific peaks and loops. ChIA-PIPE also automates quality-control assessment for each dataset. Furthermore, ChIA-PIPE generates input files for visualizing 2D contact maps with Juicebox and HiGlass, and provides a new dockerized visualization tool for high-resolution, browser-based exploration of peaks and loops. With minimal adjusting, ChIA-PIPE can also be suited for the analysis of other related chromatin-mapping data.

## Introduction

ChIA-PET (Chromatin Interaction Analysis with Paired-End Tags) enables the genome-wide discovery of chromatin interactions involving a specific protein factor (Fullwood et al., 2009). The first version ChIA-PET protocol extracted short (2 x 21 bp) paired tags for two contacting chromatin sites (Fullwood et al., 2009). Subsequently, an improved version for long read (2 x 150 bp) ChIA-PET was developed with increased mapping accuracy and robustness (Li, X. et al., 2017). ChIA-PET has been applied to study long-range chromatin interactions in human and mouse cells, and provided key insights into the roles of a number of chromatin architecture proteins and transcriptional factors including CTCF and RNAPII in human 3D-genome organization (Li et al., 2012; Dowen et al., 2014; Tang et al., 2015; Ji et al., 2016; Weintraub et al., 2017). Briefly, the long read ChIA-PET is performed as follows. First, chromatin interactions are stabilized via dual cross-linking by formaldehyde and ethylene glycol bis (succinimidyl succinate) (EGS), and then the chromatin sample is fragmented by sonication. Immunoprecipitation is performed to enrich for chromatin complexes containing a specific protein of interest. Pairs of DNA fragments in each chromatin complex are joined via bridge linker ligation (proximity ligation). The ligation products are then purified for DNA library construction and high-throughput paired-end-tag (PET) sequencing (Li X. et al., 2017).

By design, the sequencing data derived from each ChIA-PET experiment contains three sets of genomic information (Tang et al., 2015): first the genome-wide binding profile by the protein factor of interest in the study, analogous to ChIP-PET (Wei et al., 2006) and ChIP-seq data (Barski et al., 2007); second the chromatin interactions between the binding sites involving the protein factor; third, generic chromatin interactions that are not ChIP-enriched, analogous to Hi-C data (Lieberman-Aiden et al., 2009). Specifically, ChIP-enriched chromatin interactions can be distinguished based on their higher contact frequency between two given interacting loci as measured by overlapping PET counts over the non-enriched (non-specific contacts as mostly “singletons”) background.

Therefore, starting from the raw PET sequencing data of a ChIA-PET experiment, an effective computational pipeline must, as a baseline: (1) categorize read pairs (PET) for genuine bridge linker sequence in between the two genomic tags; (2) align tags to a reference genome for tag mapping locations and deduplicate the mapped reads.; (3) merge overlapping PETs to establish quantitation (PET counts) of potential chromatin contact frequency between two chromatin interaction anchor loci; (4) perform peak-calling to identify binding peaks of the protein; (5) overlap protein binding peaks with chromatin interaction anchors to identify specific-protein-involving chromatin interactions; (6) generate output of ChIA-PET data statistics and quality-assessment (QA) metrics; and (7) generate output files for data visualization and evaluation.

While computational pipelines have been developed in the past for short-reads of ChIA-PET data (Li, G. et al., 2010; Phanstiel et al., 2015) and for long-read ChIA-PET (Li, X. et al., 2017; Li, G. et al., 2017) that meet these baseline requirements, there are now significant unmet needs in the high-throughput processing and analysis of ChIA-PET data because of rapid advances in the field. First, it is now desirable to have a single pipeline with the flexibility to process either short-read (2 x 21 bp) or long-read (2 x 150 bp) data. Further, since longer read lengths increase the coverage of heterozygous SNPs, it is now desirable to have a pipeline that tests for allele-specific chromatin interactions genome-wide. Second, with falling sequencing costs, it is now common for ChIA-PET libraries to be generated with 200 - 400 million read pairs or more. An effective pipeline now requires greater robustness to process these massive data sets. Further, performing peak-calling on these larger data sets without an input-control sample, as is standard in older pipelines, results in an abundance of false-positive peaks. It is now desirable to have a pipeline that, by default, performs peak-calling with an input-control sample and with stringent parameter settings. Third, it has recently been demonstrated that chromatin contact domains (CCDs) can be effectively called from ChIA-PET data (Tang et al., 2015). Furthermore, the boundaries of CCDs called from ChIA-PET data have been shown to correspond well to the boundaries of topologically associating domains (TADs) called from Hi-C data (Tang et al., 2015). Therefore, it is now desirable to have a pipeline that automatically calls CCDs, in addition to loops and peaks. Fourth, in the past year there have been major advances in web-based visualization tools for chromatin-interaction maps with Juicebox.js (Robinson et al., 2018) and HiGlass (Kepedjiev et al., *bioRxiv*). It is now desirable to have a pipeline that generates the appropriate input visualization files (.hic file and .cool file, respectively) and that can be configured to make the files immediately visible to these tools upon completion of data processing. Fifth, variants of ChIA-PET method for studying of protein associated chromatin interactions such as HiChIP (Mumbach et al., 2016) and PLAC-Seq (Fang et al., 2016) have recently been reported. Although HiChIP data is similar to ChIA-PET data, the junction sequences between the two tags of a PET sequence are different. Therefore, it has now desirable to have a pipeline that is capable of processing both ChIA-PET and HiChIP data.

ChIA-PIPE is designed and integral with multiple components to achieve all of these emerging needs. The pipeline is seamlessly integrated and runs from a single launch command, while also having the modularity to be re-entered at any step (Figure 1, Figure S1). The pipeline is parallelized, open source, and can be applied to a new ChIA-PET library simply by modifying a configuration file. In addition, ChIA-PIPE generate output files for data visualization in 2D contact maps (JuiceBox and HiGlass) and browser-based views. More specifically, we introduce a new loop browser, called BASIC Browser, for interactive, high-resolution visualization of loops, domains, and binding coverage. ChIA-PIPE is currently used for all ChIA-PET data processing as part of the ENCODE4 and 4D Nucleome consortia.

**Figure 1.**
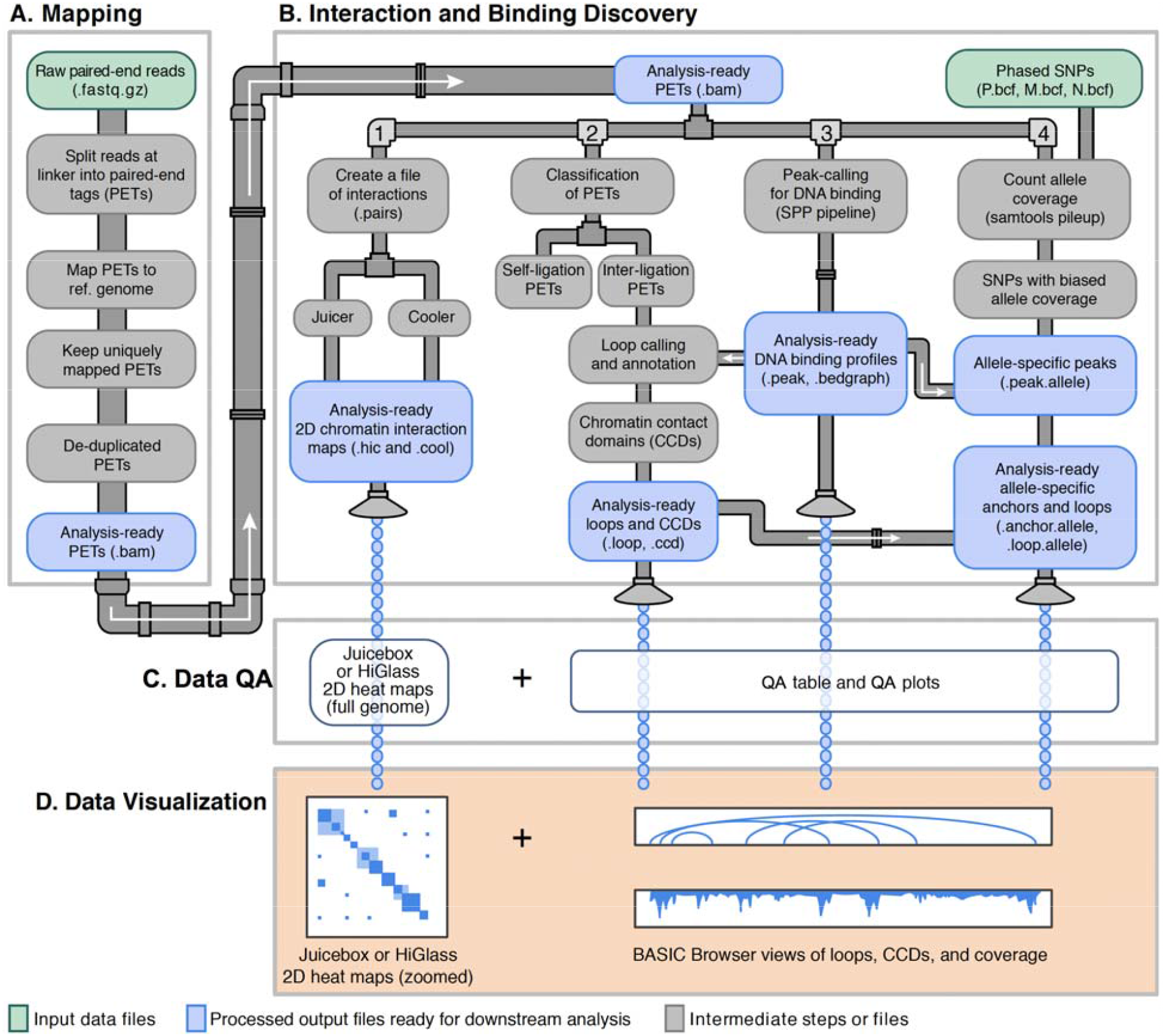
ChIA-PIPE architecture. ChIA-PIPE takes as input a configuration file and two FASTQ files (of R1 reads and R2 reads). (A) First, read pairs are scanned for the bridge-linker sequence and partitioned into categories: read pairs with (i) no linker, (ii) a linker and a one usable genomic tag, or (iii) a linker and paired-end tags (PETs). PETs are aligned to a reference genome. The analysis-ready BAM file contains uniquely-mapped, non-redundant PETs. (B.1) 2D chromatin-interaction maps are generated with standardized file formats. (B.2) Using inter-ligation PETs (span ≥ 8 kilobases), loops are called and then annotated with peak support. Peak-supported loops are then used to call chromatin contact domains (CCDs). (B.3) Binding peaks of the protein factor are identified using SPP (Kharchenko et al., 2008) (or optionally, MACS2 (Zhang et al., 2008)). Prior to peak calling there is a recovery step to incorporate all uniquely-mapped, non-redundant tags (not only linker containing PETs). (B.4) If phased SNP information is available for the cell type of interest, allele-specific peaks and loops are identified. First, SNPs with significant bias in allele coverage are identified. Then, biased SNPs are used to annotate peaks and loop anchors. (C) The pipeline collates quality assessment (QA) metrics from every step into a succinct QA table and generates extensive QA plots. (D) The output files from the pipeline are compatible with interactive, high-resolution visualization tools. The 2D chomatin-interaction maps can be viewed in Juicebox.js (Robinson et al., 2018) or HiGlass (Kerpedjiev et al., bioRxiv). The loops file, CCD file, and coverage file can be viewed in BASIC Browser.

## Results and Discussion

The ChIA-PIPE contains four modules (Figure 1): A. Mapping; B. Interaction and binding discovery, C. data quality assessment, and D. data visualization. For massive data sets with, almost every step of the pipeline supports multi-threading. Also, the config file allows the user to customize the number of threads and the memory. The input for ChIA-PIPE is a configuration file and two FASTQ files (one file for R1 reads and one file for R2 reads). The first step (Figure 1A) is to scan the read pairs for the linker sequence from proximity-ligation step of ChIA-PET and to partition the read pairs into categories: read pairs with (i) no linker sequence, (ii) a linker sequence and one usable genomic tag, or (iii) a linker sequence and paired-end tags (PETs). ChIA-PIPE also supports the processing of HiChIP data, by treating the repaired and ligated restriction site from HiChIP as a “pseudo-linker” in this first step (see Methods).

Next, each category of read pairs is separately aligned to a reference genome (see Methods). Only uniquely mapped and non-redundant tags are retained in each BAM file. The BAM file of PETs is the final output file then used for all downstream analyses of chromatin interactions (Figure 1A). Only the peak-calling step uses a merge of all three BAM files (as even tags that were non-interactions are still informative for binding).

Once the BAM file of analysis-ready PETs is generated, ChIA-PIPE performs interaction and binding discovery in several workflows. First, the BAM is converted into a standardized file format for interaction pairs, which is then converted into 2D contact-map files for visualization of chromatin-interaction heat maps using Juicebox.js or HiGlass (Figure 1B.1).

ChIA-PIPE also identifies genomic binding peaks of the protein factor (Figure 1B.3). By default, this peak-calling is performed using SPP (Kharchenko et al., 2008), which identifies genomic regions of significantly enriched tag density compared to a standard ChIP-seq input-control sample. In practice, this approach has exhibited high specificity even with large data sets, and has become an ENCODE standard. ChIA-PIPE also has the flexibility for peak calling to be optionally performed using MACS2 (Zhang et al., 2008) with or without a ChIP-seq input-control sample.

In addition, ChIA-PIPE calls and annotates loops, as well as calling chromatin contact domains (CCDs) (Figure 1B.2). Using “inter-ligation” PETs, each tag is extended by 500 bp in its 5’ direction and overlapping PETs are merged into “loops”, where the intensity of the loop is measured by the count of the PETs contributing to the loop (see Methods). Each loop is then annotated with the number of its anchors (0, 1, or 2) that are supported by a binding peak. ChIA-PIPE provides the chromatin interaction and peak-annotated anchors file for users to further apply existing tools (Paulsen et al., 2014; He et al., 2015; Phanstiel et al., 2015) to identify significant loops. CCDs are then called using loops with peak support (see Methods). If phased SNP data is available for the appropriate cell type, ChIA-PIPE identifies peaks and loops with significant allele bias (Figure 1B.4).

To enable rapid evaluation of library quality, the pipeline reports key data statistics and quality assessment metrics in a CSV file (Figure 1C and Figure S2), which can be easily opened as an Excel spreadsheet. For example, the majority of the read pairs should contain the linker sequence and, of those, the majority should be PETs (rather than one-tag read pairs). Further, ChIA-PIPE supports QA visualization for each library. Library-level QA visualizations reveal valuable information about: the percentage of read pairs passing each processing step, the quality of the binding enrichment, and the specificity of the chromatin interactions (Figure 2A-C).

**Figure 2.**
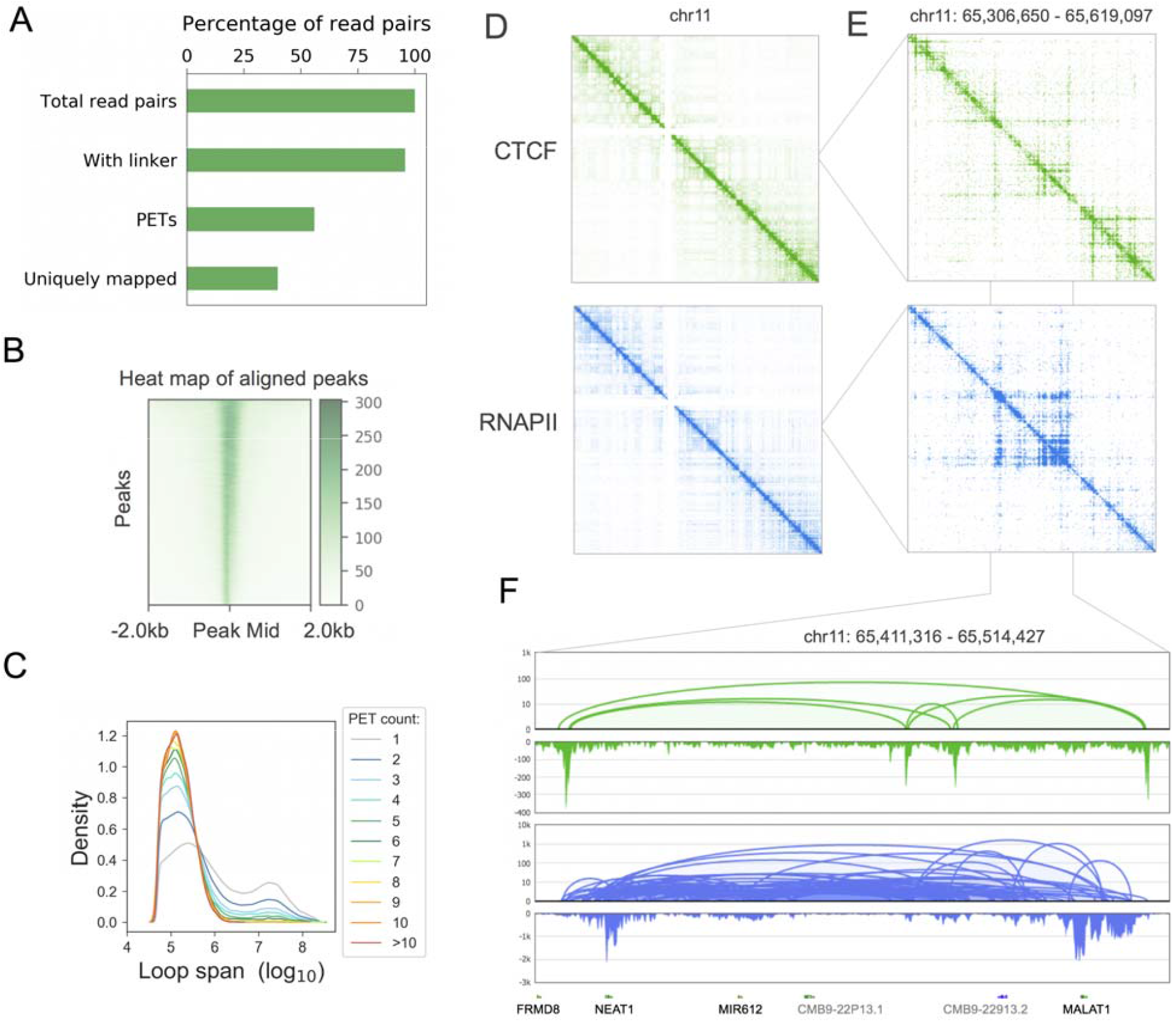
ChIA-PIPE quality assessment and visualization. ChIA-PIPE supports quality assessment (QA) and visualization for each sequencing library that is processed. (A) A breakdown of the percentage of read pairs passing various data pre-processing steps (for a Hiseq CTCF ChIA-PET library in HFFc6 cells). The pipeline also generates a QA table with more detailed summary statistics. (B) A peak intensity tornado heatmap of the read pileup over all peaks (each row) aligned by their midpoints (shown: CTCF in HFFc6). The sharpness indicates the quality of the binding enrichment). (C) The distributions of loop span depending on the PET count of the loops (shown: CTCF in HFFc6). (D-E) Juicebox.js can be used for QA visualization by viewing the 2D contact map for full chromosome or broad chromosomal segment. The enrichment of signal along the diagonal indicates the quality of the library. As examples, we showed both CTCF (D) and RNAPII (E) ChIA-PET data in chr11 and a genomic region (chr11:65,306,650 – 65,619,097). (F) BASIC browser provides detailed and high-resolution visualization of smaller chromosomal regions. BASIC browser can be used for QA visualization by viewing a known biological interaction (such as NEAT1 and MALAT1) to confirm looping and binding enrichment (shown: CTCF and RNAPII in HFFc6).

Notably, ChIA-PIPE automatically generates 2D contact-map files for both web-based tools: a .hic file for Juicebox.js (Robinson et al., 2018) and a .cool file for HiGlass (Kepedjiev et al., *bioRxiv*). The 2D contact maps provide informative views of full chromosomes or moderately sized chromosomal regions (Figure S2B-C, 2D-E). BASIC Browser provides detailed and high-resolution views of even smaller chromosomal regions (Figure 2F).

The ENCODE4 and 4D Nucleome consortia are generating thousands of ChIA-PET libraries over the next few years - each library contains 200-400 million next generation sequencing read pairs. ChIA-PIPE is designed to robustly to process the massive datasets by parallelizing with multiple threads, when appropriate hardware is available. For example, ChIA-PIPE processed HFFc6 CTCF HiSeq library (Figure 4A, ~300 million read-pairs) follow the procedures described above with run time (wall) equal to 10 hours (Threads = 20, RAM = 60 GB, Cluster OS = CentOS 6.5, CPU = Intel Xeon E5-2670 @ 2.60GHz).

In case where phased SNP data is available from matched cell type for the ChIA-PET data, one can also derive allele-specific peaks and loops using ChIA-PIPE. In Figure 3A, we show the detailed ChIA-PIPE allele-specific peak and loop calling workflow: first, SNPs with significant allele bias in the coverage of ChIA-PET reads are identified (see Methods); second, peaks overlapping significantly biased SNPs are kept based on the results of binomial test (see Methods); third, loop anchors overlapping significantly biased peaks are identified (see Methods). The ChIA-PIPE allele-specific peak and loop output report includes the paternal and maternal biased SNPs, peaks, and loops (Figure 3B).

**Figure 3.**
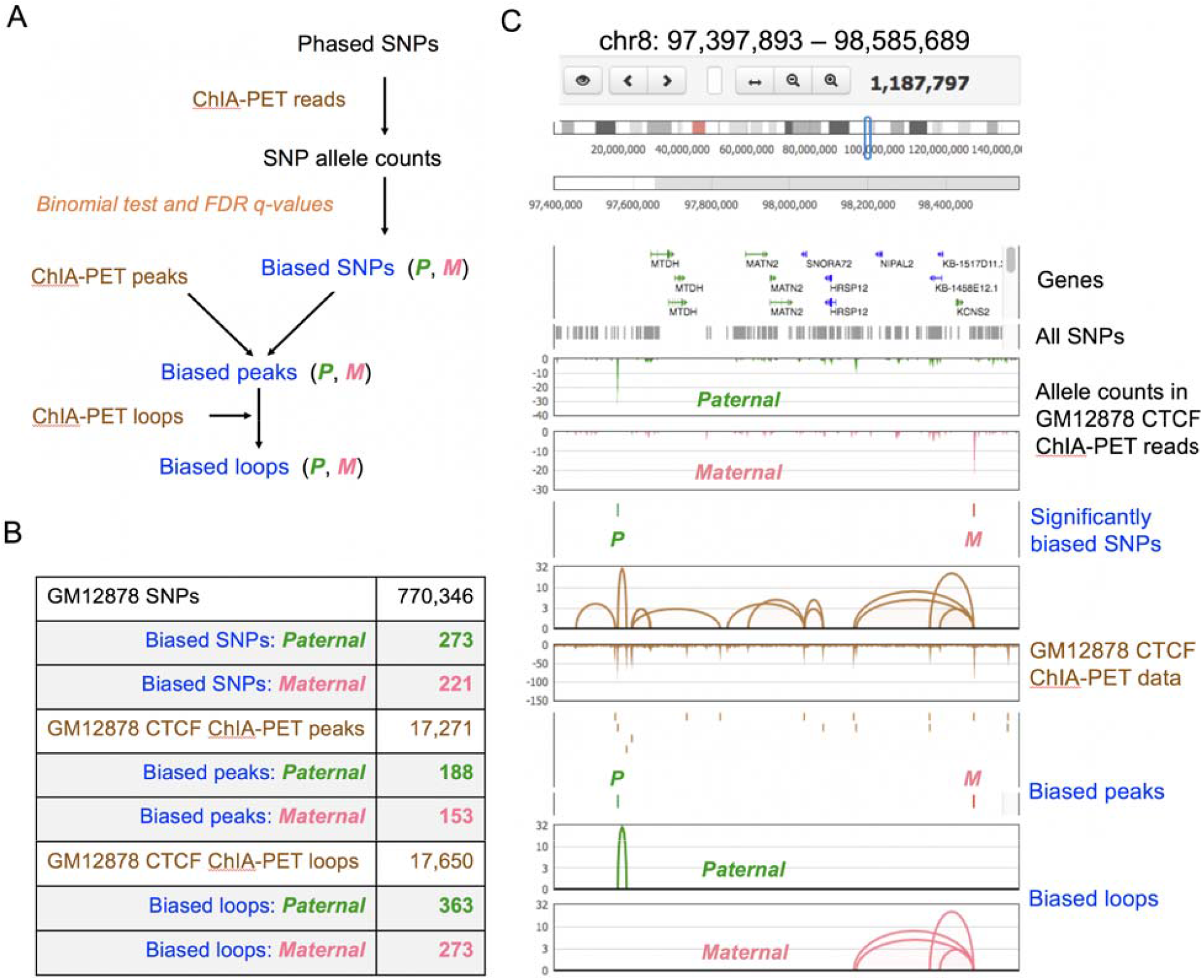
ChIA-PIPE can resolve allele-specific chromatin interactions. If phased SNP information is available for the cell type of interest, ChIA-PIPE can resolve allele-specific peaks and loops. (A) First, for each phased SNP, the alleles counts in the ChIA-PET reads are determined. Then, SNPs with significantly biased allele counts are determined. ChIA-PET peaks that overlap significantly biased SNPs are then considered biased peaks. ChIA-PET loops that overlap biased peaks and then considered biased loops. (B) An example of the maternal and paternal number of biased SNPs, peaks, and loops from GM12878 CTCF ChIA-PET MiSeq data. (C) An example visualization in BASIC Browser of a genomic region with one paternally biased SNP and one maternally biased SNP, and the corresponding biased ChIA-PET peaks and loops.

Once a library has been processed and passes QA, gaining insight into the novel biological findings in the library requires high-resolution, interactive visualization of ChIA-PET data in browsers. The pipeline automatically outputs the appropriate file formats for all relevant 1D- and 2D-interactive visualization tools for ChIA-PET data (Figure 1D). For 1D visualization, the key tracks for ChIA-PET data are the loop clusters (a BEDPE file), the CCDs (a BED file), and the binding-intensity profile of the protein factor (a bedgraph file). The binding-intensity track can be readily viewed in two publicly-available genome browsers: UCSC Genome Browser (Rosenbloom et al., 2013) and the WashU Epigenome Browser (Zhou et al., 2013). However, only the latter can readily display chromatin-interaction loops, and even then, the display is not the most intuitive, as the loop height (y axis) does not scale with the PET count.

Therefore, ChIA-PIPE includes its own loop browser -- “Browser for Applications in Sequencing and Integrated Comparisons” (BASIC Browser) -- for interactive, high-resolution ChIA-PET data visualization (Figure 3C and Figure 4). In BASIC browser, the chromatin-interaction tracks display the chromatin loops for their contact frequency (y-axis, the height of a loop reflects the intensity of connectivity) and interacting anchor distances, along with the protein binding-intensity tracks. The BASIC browser tracks are intuitive and in publication-quality ready, which can reveal novel biological findings (Figure 3C, Figure 4, Figure S4). For ease of use, BASIC Browser is made available as a Docker image.

**Figure 4.**
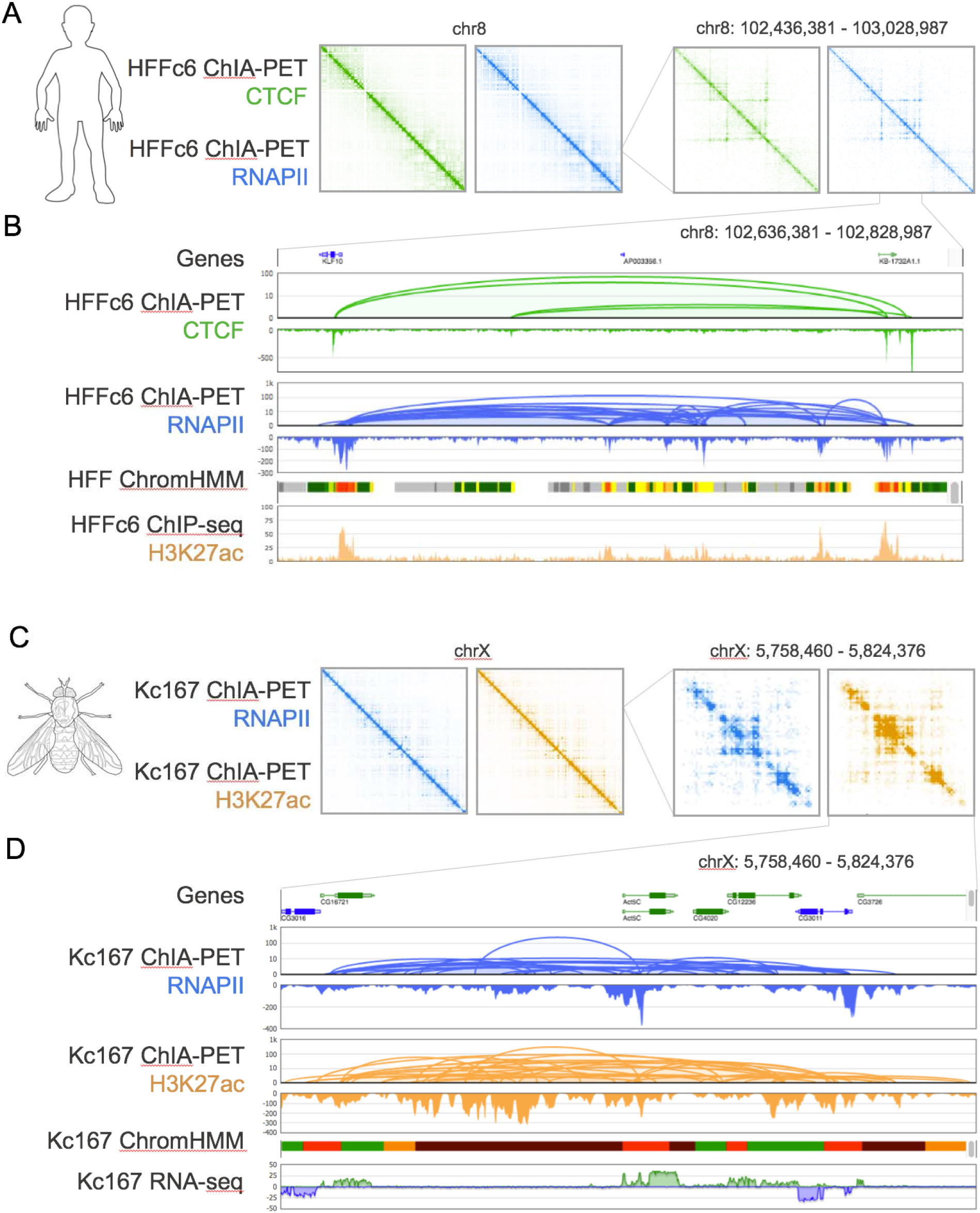
ChIA-PIPE enables visualization of 2D contact maps with Juicebox and HiGlass, and browser-based views of loops and peaks. ChIA-PIPE includes a Docker image of BASIC Browser for interactive, high-resolution ChIA-PET data visualization. (A) Juicebox heat maps of CTCF and RNAPII ChIA-PET, showing full chromosome 8 and then a broad region on chromosome 8. (B) BASIC browser provides detailed and high-resolution visualization of smaller region on chromosome 8. Shown are the loops, chromatin contact domains (CCDs), and binding coverage for CTCF ChIA-PET and RNAPII ChIA-PET in HFFc6 cells. In addition, BASIC Browser supports the visualization of other data tracks to facilitate ChIA-PET interpretation. For example, the UCSC Known Genes are shown at the top and ChromHMM in the same cell type (from the ENCODE portal) and H3K27ac ChIP-seq in the same cell type (from the 4DN) portal are shown at the bottom. For the ChromHMM track, red indicates transcription start sites (TSSs) and yellow and orange indicate enhancers. Thus, BASIC browser provides a comprehensive genomic view of this region, revealing that a CTCF-mediated loop is encompassing two active genes that are interacting with each other and with a set of enhancers between them. (C) Juicebox heatmaps of CTCF and RNAPII ChIA-PET from *Drosophila* Kc167 cells. (D) A more detailed view of the data in BASIC browser, also displaying the UCSC Known Genes, the ChromHMM track in the same cell type, and RNA-seq in the same cell type.

In summary, ChIA-PIPE has achieved multiple milestones: (**1**) full automation as a single launch command-based pipeline; (**2**) robustness to massive dataset (**3**) ChIA-PET sequencing read length flexibility; (**4**) accurate peak calling achieved by SPP with an input control sample; (**5**) automated chromatin contact domain (CCD) calling; (**6**) web-based visualization automation for BASIC browser, Juicebox.js and Higlass; and (**7**) adaptability to process data from related protocol (e.g., HiChIP). These achievements have made ChIA-PIPE a valuable resource used by the ENCODE4 and 4D Nucleome consortia, and the broader biological research community.

## Methods

### Linker filtering

The linker-filtering step employs ChIA-PET Utilities (CPU, https://github.com/cheehongsg/CPU), a collection of modularized executables developed by our team for performing core ChIA-PET data-processing tasks. ChIA-PIPE integrates CPU modules and modules from many other packages into a comprehensive pipeline (Figure S1). Though HiChIP data does not contain a linker sequence, all read pairs with interactions contain a repaired and ligated restriction site (GATCGATC), which can be treated as a “pseudo-linker” in the pipeline.

### Tag alignment

Tag alignment uses the CPU module “memaln” a combination of bwa mem and bwa aln (Li H. et al., 2009a), which can handle either short-read or long-read data.

The input file of the ChIA-PET sequencing data is fastq.gz. Briefly, paired-end tag (PET) read sequences were scanned for the bridge linker sequence and only PETs with the bridge linker were retained for downstream processing. After trimming the linkers, the flanking sequences were mapped to the human reference genome (hg19) using a hybrid of BWA-MEM and BWA-ALN, and only uniquely aligned (MAPQ ≥ 30) PETs were retained. PCR duplicates were removed using the MarkDuplicates tool of the Picard Tools library (http://broadinstitute.github.io/picard/). The BAM file of PET reads is now ready for further analysis (Figure 1A).

### Interaction and binding discovery

The BAM file (.bam) of PET reads alignment is further processed in 4 parallel sub-pipelines for identifications of chromatin interaction, 1) 2D contact maps, 2) loops, 3) binding peaks, and 4) haplotype-specific interactions.

1. *2D contact maps* (Figure 1B.1). The BAM file of all analysis-ready PETs is processed to generate 2D contact matrix file (.hic) using Juicer tools (Durand, N.C., et al., 2016a). We generate the contact matrix file for 10 different resolutions (2.5Mb, 1Mb, 500Kb, 250Kb, 100Kb, 50Kb, 25Kb, 10Kb, 5Kb, 1Kb), and thus users can easily adjust multi-scale genomic regions from all chromosomes view to a specific genomic region, up to 1Kb resolution. Juicebox, described at the data visualization section, is used to access the 2D contact map (Durand, N.C., et al., 2016b). Based on the 4DN DCIC’s request, the BAM file of all analysis-ready PETs will also be processed to generate a contact list file (.pairs) that is compatible with the proposed 4DN DCIC’s pipeline for Hi-C data analysis.
2. *Interaction loops* (Figure 1B.2). This step of the pipeline employs the CPU module “cluster”. The BAM file of PET reads are from three categories: (a) PET reads with no linker sequence detected, (b) PET reads with a linker sequence detected but with only one usable genomic tag and (c) PET reads with a linker sequence detected with both ends having genomic tags. Only PET reads in (c) are used for further detecting interaction loops. Each PET in (c) was categorized as either a self-ligation PET (two ends of the same DNA fragment) or inter-ligation PET (two ends from two different DNA fragments in the same chromatin complex) by evaluating the genomic span between the two ends of a PET. PETs with a genomic span less than or equal to 8 kb are classified as self-ligation PETs and are used as a proxy for ChIP fragments since they are derived in a manner analogous to ChIP-Seq mapping for protein binding sites. PETs with a genomic span greater than 8 kb are classified as inter-ligation PETs and represent the long-range interactions of interest. To be more representative of the interacting chromatin fragments, the 5’ end of each inter-ligation PET was extended by 500 bp along the reference genome. To reflect the frequency of interaction between two loci, the extended inter-ligation PETs that directly overlap were clustered together as one PET cluster. The PET counts in a PET cluster reflects the relative frequency of interaction between two genomic regions. It was observed that many anchors of distinct PET clusters were located within the same protein factor binding peak. It is clear that these binding peaks reflect the real chromatin interaction loci in the nucleus. In order to streamline the PET clusters data structure, we collapsed the individual anchors of all PET clusters with 500 bp extensions to generate merged anchors. For anchors with overlapped binding peaks, we use the summit point as the centers of interacting loci. We refer the merged PET clusters as chromatin interaction loops. Un-clustered individual inter-ligation PETs and PETs in the clusters below the PET cutoff are referred as PET singletons.
3. *Interaction binding peaks* (Figure 1B.3). All uniquely mapped and non-redundant analysis-ready reads including self-ligation and inter-ligation are used for identifying protein factor binding peaks. In addition, two categories of PET reads that were excluded from chromatin loop detection are recovered and included in peak-calling for protein factor binding. Specifically, the recovered read categories are (a) PET reads with no linker sequence detected and (b) PET reads with a linker sequence detected but with only one usable genomic tag. While these reads are uninformative for chromatin loop detection, they can be informative for peak-calling of protein factor binding. Peak calling for protein factor binding is then performed using the SPP pipeline (version 1.13) (Kharchenko et al., 2008) with parameters srange=c(200,5000), bin=20, window.size=500 and z.thr=6. Optionally, the MACS2 pipeline (version 2.1.0) (Zhang et al., 2008) can be used for protein factor binding peak identification with default parameters. In addition, bedtools is used to generate BedGraph files of the protein factor binding coverage along the chromosomes for browser-based visualization.
4. *haplotype-specific interactions* (Figure 1B.4 and Figure 3A). There are several steps involved in determining haplotype-specific chromatin interactions, if the phased genome sequencing information is available. Using GM12878 cell line as an example, it is derived from the 1000 Genome Project human subject NA12878, whose genome-wide single nucleotide polymorphism (SNP) phasing information is available (ftp://gsapubftp-anonymous@ftp.broadinstitute.org/bundle/2.8/hg19/). By using this SNP phasing information, uniquely aligned (MAPQ ≥ 30) CTCF and RNAPII ChIA-PET reads were given a haplotype assignment depending on whether or not they overlapped a phased SNP and, if so, which allele they had. The possible haplotype assignments were maternal (M), paternal (P), or not determined (N). This analysis was done independently for the two ends of all inter-ligation PETs. Therefore, a PET could have the possible paired haplotype at the two ends of intra-chromosomal loops M-M, P-P, M-N, P-N, M-P, or N-N. The allele counts were determined using samtools pileup (Li H. et al., 2009b).

Before this was done, the haplotype specificity of each interaction anchor was determined. The haplotype assignment of the anchors was performed as follows: 1) Phased SNPs with biased protein factor binding coverage were identified. The maternal and paternal allele counts of individual phased SNPs were computed and tested for allele bias using a Binomial test. SNPs with Benjamini-Hochberg-adjusted P-values (i.e., FDR Q-values) ≤ 0.1 (Yoav, B. et al., 1995). were considered to have significantly biased protein factor binding coverage; 2) The interaction anchors that overlap with biased SNPs were then given a haplotype assignment corresponding to the bias direction of the SNP. If an anchor overlapped with multiple biased SNPs and the bias directions of these SNPs were consistent, then the haplotype assignment of this anchor is given accordingly; otherwise, the haplotype of this anchor was considered to be not determined. In addition, if an anchor overlapped with multiple phased SNPs and the SNP with the highest binding coverage showed no allelic bias, then the haplotype of such an anchor was also considered to be not determined. The above procedures were performed in the CTCF and RNAPII ChIA-PET datasets independently. For CTCF ChIA-PET MiSeq data from GM12878 cells, 636 CTCF interaction loops were identified as Phased Interactions. Allele-specific chromatin loops and anchors can also be used to show chromatin loop specificity between cells with different genetic backgrounds (Figure 3 and S3).

### ChIA-PET QA visualizations

For each ChIA-PET library, we first perform a QA test sequencing to generate 5-10 million PET reads from a MiSeq run, which usually can test 2-4 ChIA-PET libraries. Once a quality library is verified by the test sequencing data, we then generate ~200 million PET reads from a HiSeq “production” run. Both of the test and production sequencing data are processed using the ChIA-PET data processing pipeline detailed below. Currently the pipeline provides library level quality examination: sequencing alignment quality, peak intensity tornado plot (generated by deepTools2, Ramírez F. et al., 2016), loop spanning distribution, and loop level anchor support distribution. The HiC file generated by the pipeline can be ready to examined by Juicebox.

### Interactive, high-resolution ChIA-PET data visualization

ChIA-PET mapping data can be viewed in 1-D browser tracks for protein binding peaks, chromatin loops between the interaction anchors of binding peaks, and the 2-D heat maps of chromatin contacts.

1. *BASIC Browser for interactive, high-resolution visualization of ChIA-PET loops, domains, and binding coverage*. We use an in-house developed genome graphic browser to visualize the binding peaks and chromatin loops (Figure 4). The chromatin interaction data file (loop.gz; Figure 1B.2) is uploaded to the ChIA-PET browser and display chromatin contacts as arcs between each of the paired genomic loci. The length of an arc indicates the linear genomic distance between two connecting loci, and the height of an arc reflects the contact frequency (PET counts) detected in the ChIA-PET data. The files for protein binding profiles (Figure 1B.3) are uploaded to the ChIA-PET browser to visualize the binding peaks and to demarcate the summit of binding sites. In addition, haplotype-specific files for binding peaks and chromatin interactions (Figure 1B.4) are also visualized. The advantages of using browser-based visualization includes detailed base-pair resolution presentation of chromatin loops in relation to protein peaks, and readily to be integrated with other genome browser information of other experimental data (RNA-Seq, ChIP-Seq, ATAC-Seq, etc).
2. *Interactive, high-resolution visualization of ChIA-PET 2D contact maps using Juicebox.js and HiGlass*. We use Juicebox.js and HiGlass to visualize ChIA-PET data in 2-D heat maps. After the duplications are removed and the uniquely mapped PETs are retained, the bam file can be converted to a merged_nodups.txt file and used as input to the Juicer tools Pre command, creating a .hic file. Loops called by ChIA-PET software can also be visualized in Juicebox by representing them in the appropriate text file format. Juicer is a data processing tool to generate the contact matrix file (.hic), a highly compressed binary file format, and the contact matrix file can be accessed by Juicebox (Durand, N.C., et al., 2016b), visualization software. These tools were developed by Aiden’s Lab to initially visualize HiC data, and we adopt these tools to process and visualize our ChIA-PET 2D contact heat maps. We generate the contact matrix file for 10 different resolutions (2.5Mb, 1Mb, 500Kb, 250Kb, 100Kb, 50Kb, 25Kb, 10Kb, 5Kb, 1Kb) using Juicer tool, and thus users can easily adjust multi-scale genomic regions from all chromosomes view to a specific genomic region, up to 1Kb resolution. Contact matrix is based on the adjustable resolution, from 2.5Mb x 2.5Mb bin size to 1Kb x 1Kb bin size. Raw data, or none normalization data, will be the total contact count within the specific genomic bin. Contact signal intensity, or counts, represent by red color, from low contact (dim red) to high contact (dark red). Four different normalization methods (none, coverage, coverage_sqrt, balanced) can be applied to the contact heat map data to remove technical noise or adjust biased signal.

## Declaration

### Availability of software

ChIA-PIPE is implemented in bash, awk, Python, Perl, and R and is freely available, along with documentation, at https://github.com/TheJacksonLaboratory/ChIA-PIPE. Source code is additionally included as Supplemental Material. BASIC Browser is available as Docker image, which is freely available at https://github.com/TheJacksonLaboratory/basic-browser.

### Availability of data and materials

The HFFc6 ChIA-PET data used in the current paper have been deposited to 4D Nucleome Portal, and are available for download at https://data.4dnucleome.org/experiment-set-replicates/4DNESCQ7ZD21/. The GM12878 HiChIP data used in this manuscript was downloaded from the Sequence Read Archive (SRR3467175).

## Acknowledgements

Y.R. was supported by ENCODE grant UM1 HG009409, 4D Nucleome grant (U54 DK107967), and JAX Director’s Innovation Award Director’s Innovation fund 19000-18-02. S.L. was supported by Leukemia Research Foundation New Investigator Grant, The Jackson Laboratory Director’s Innovation fund 19000-17-31, and The Jackson Laboratory Cancer Center New Investigator Award. Research reported in this publication was partially supported by the National Cancer Institute of the National Institutes of Health under Award Number P30CA034196. The content is solely the responsibility of the authors and does not necessarily represent the official views of the National Institutes of Health. The authors thank members of the Wei Lab, Ruan Lab, Li Lab, Chee-Hong Wong, Joshy George, and Idan Gabdank for helpful discussions, thank Zoe Reifsnyder for artistic improvement of the figures, and thank the legacy programmers who contributed to BASIC browser during its early development.

---

**Figure S1.**
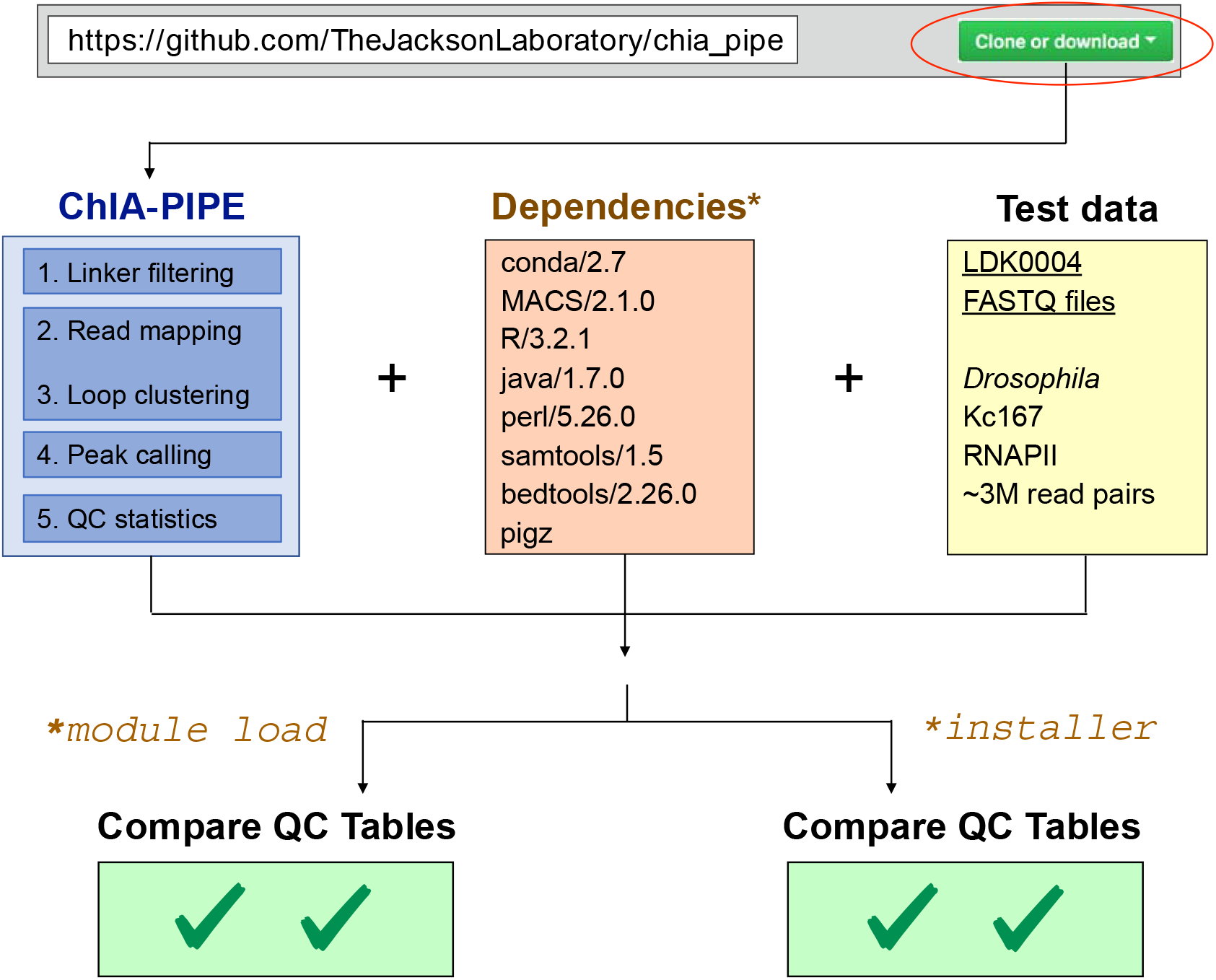
ChIA PIPE installation. The ChIA PIPE source code can be downloaded from its Github repository. The dependencies of ChIA PIPE are listed in the orange box in the middle. In some high-performance computing clusters, the dependences can simply be loaded using the ‘module load’ command. Otherwise, ChIA-PIPE also includes an installation script for performing a local install of the dependencies. Finally, a small test set is provided of RNAPII ChIA-PET in *Drosophila* Kc167 cells. When this data set is processed, the resulting quality-assessment table can be compared to a reference quality-assessment table to ensure proper installation of the pipeline.

**Figure S2.**
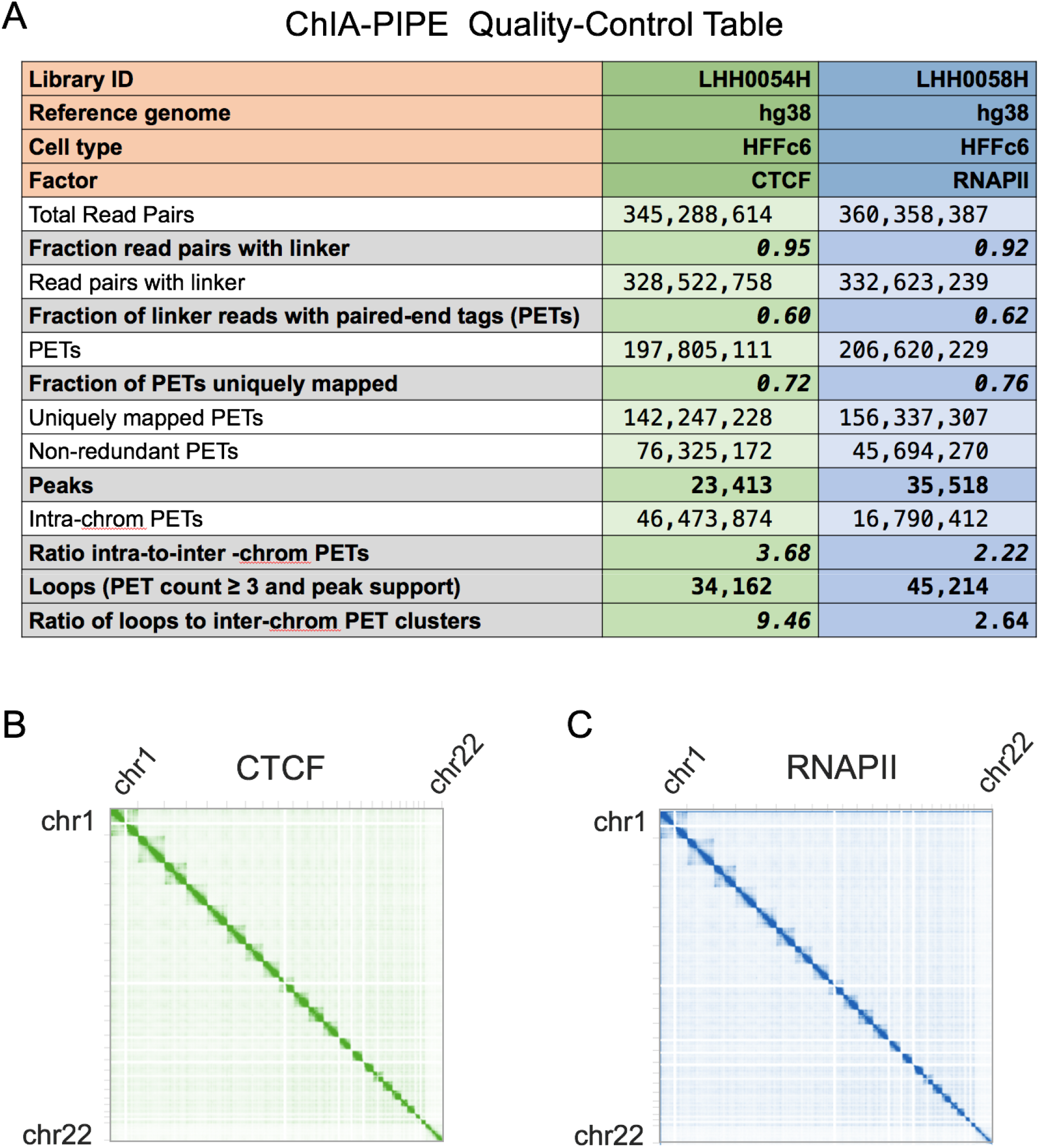
ChIA PIPE quality-assessment visualization. (A) An example quality assessment (QA) table generated by ChIA-PIPE. QA results are shown for one CTCF library and one RNAPII library in HFFc6 cells. Key parameters are: the fraction of read pairs with linkers (indicating ligation efficiency), the fraction of paired-end tags (indicating tagmentation optimization), the fraction of PETs uniquely mapped (indicating the amount of usable data), the number of called binding peaks, the number of loops, and the ratio of intra-to-inter chromosomal PETs and the ratio of intra-to-inter chromosomal loops (both indicating the signal to noise ratio of the data). (B-C) Full-genome Juicebox heat maps are shown for CTCF ChIA-PET (B) and RNAPII (C) ChIA-PET in HFFc6 cells (indicating the signal to noise ratio of the data).

**Figure S3.**
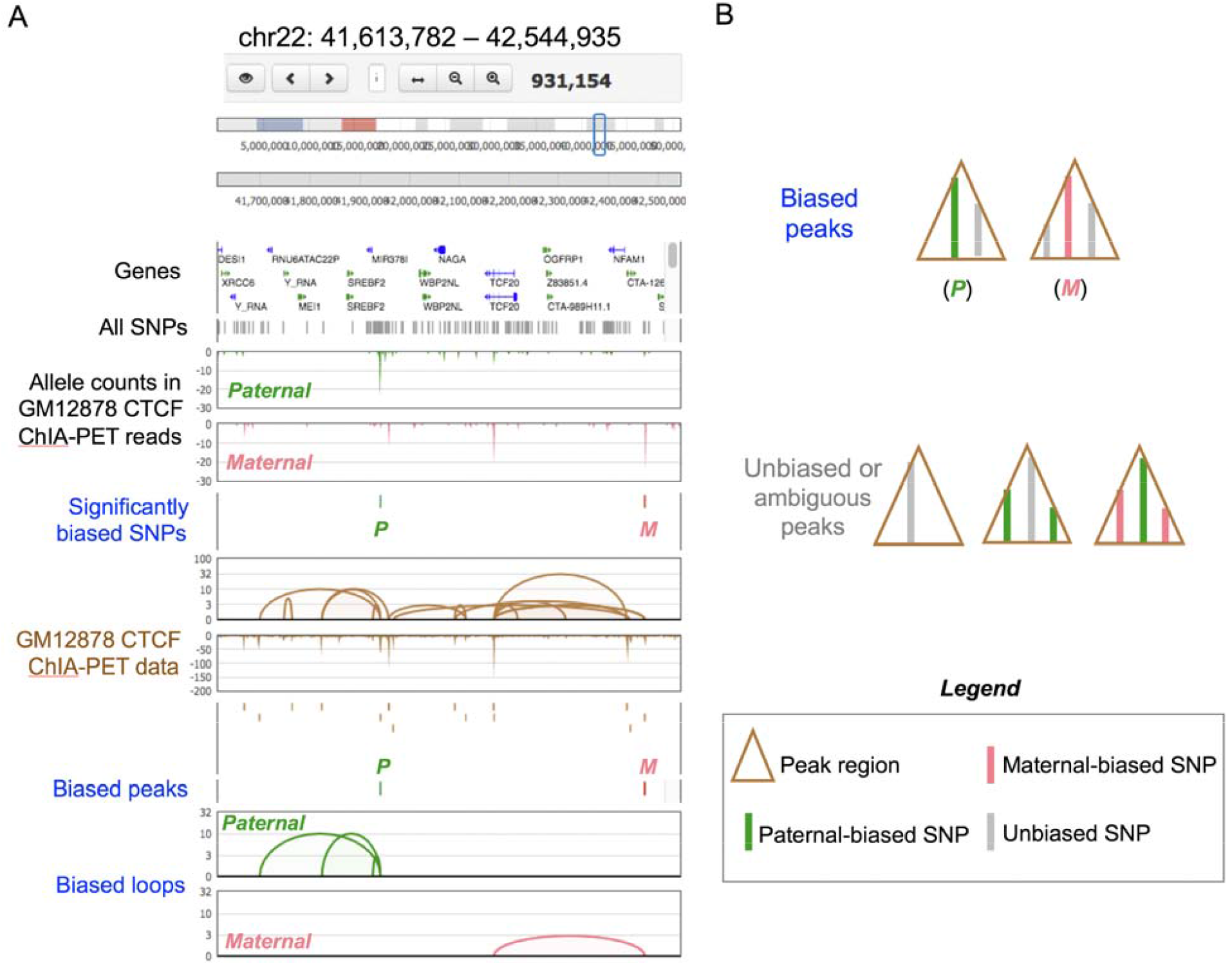
ChIA-PIPE resolves allele-specific chromatin interactions. (A) An additional BASIC Browser example of a genomic region with one paternally biased SNP and one maternally biased SNP, and the corresponding biased ChIA-PET peaks and loops. (B) A diagram demonstrating how ChIA-PIPE categorizes peaks that overlap more than one SNP. In such cases, a peak is considered biased only if the SNPs are biased in the same direction (paternally biased SNPs and unbiased SNPs; or maternally biased SNPs and unbiased SNPs) and if the SNP with the highest read coverage in the peak is a biased SNP.

**Figure S4.**
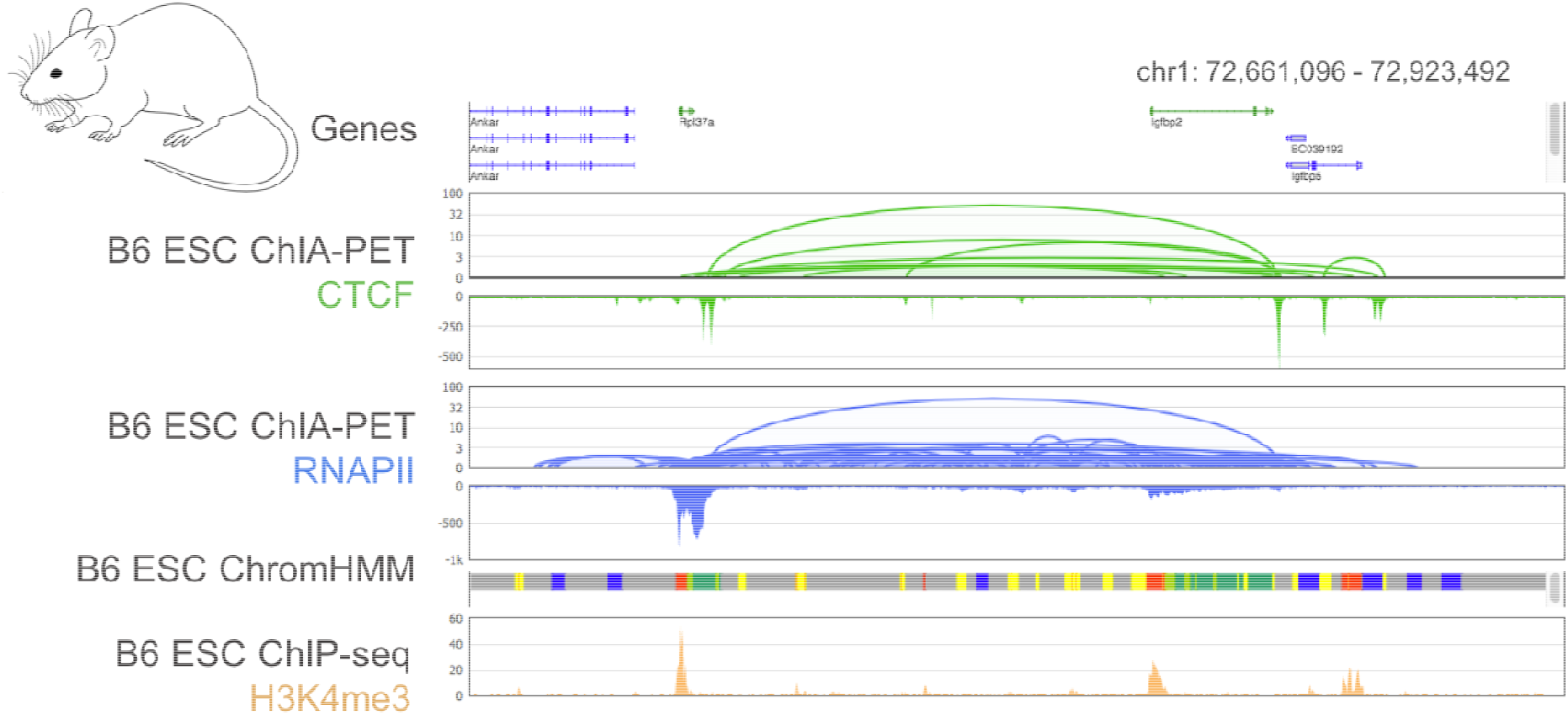
BASIC Browser for interactive, high-resolution visualization of ChIA-PET loops, domains, and coverage. An additional BASIC Browser example of CTCF and RNAPII ChIA-PET data in mouse embryonic stem cells. Shown are the ChIA-PET loops and binding coverage. BASIC Browser supports the visualization of other data tracks to facilitate ChIA-PET interpretation. For example, the UCSC Known Genes are shown at the top and ChromHMM in the same cell type (from the ENCODE portal) and H3K4me3 ChIP-seq in the same cell type (from the 4DN) portal are shown at the bottom.

**Figure S5.**
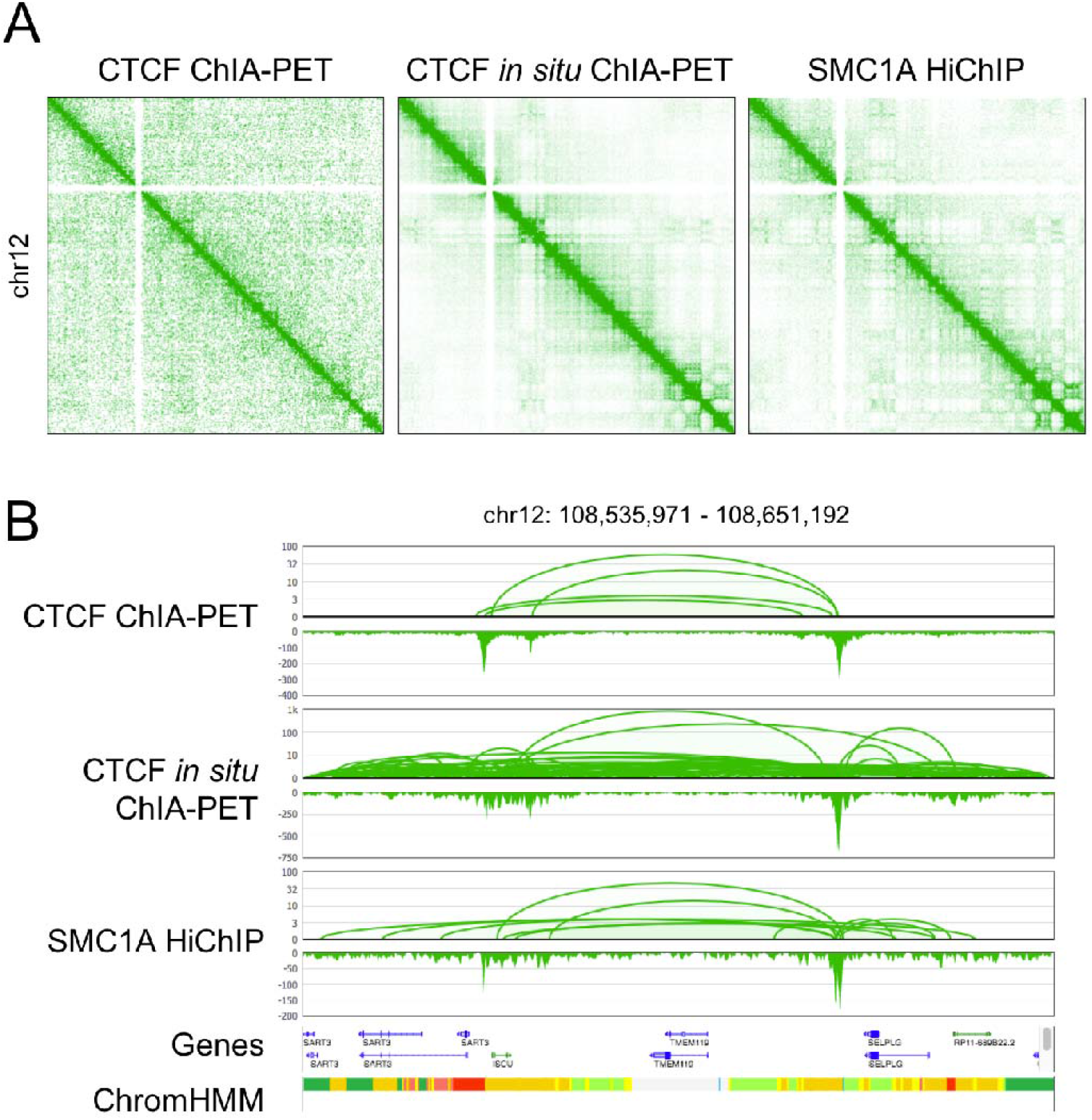
ChIA-PIPE can be used to process data from related 3D-genome mapping methods, such as HiChIP. ChIA-PIPE was used to process comparable data sets in GM12878 cells from different methods: CTCF ChIA-PET, CTCF *in situ* ChIA-PET, and SMC1A (cohesin) HiChIP. ChIA-PIPE was readily adapted to process HiChIP data by treating the ligated restriction enzyme site as a “pseudo-linker”. (A) 2D heat maps of the three data sets from Juicebox.js. (B) BASIC Browser views of loops and binding coverage from the three data sets, alongside gene annotations and ChromHMM in GM12878 (from the ENCODE portal).

