## Supplemental Figures for "ChIA-PIPE: A fully automated pipeline for ChIA-PET data analysis and visualization"

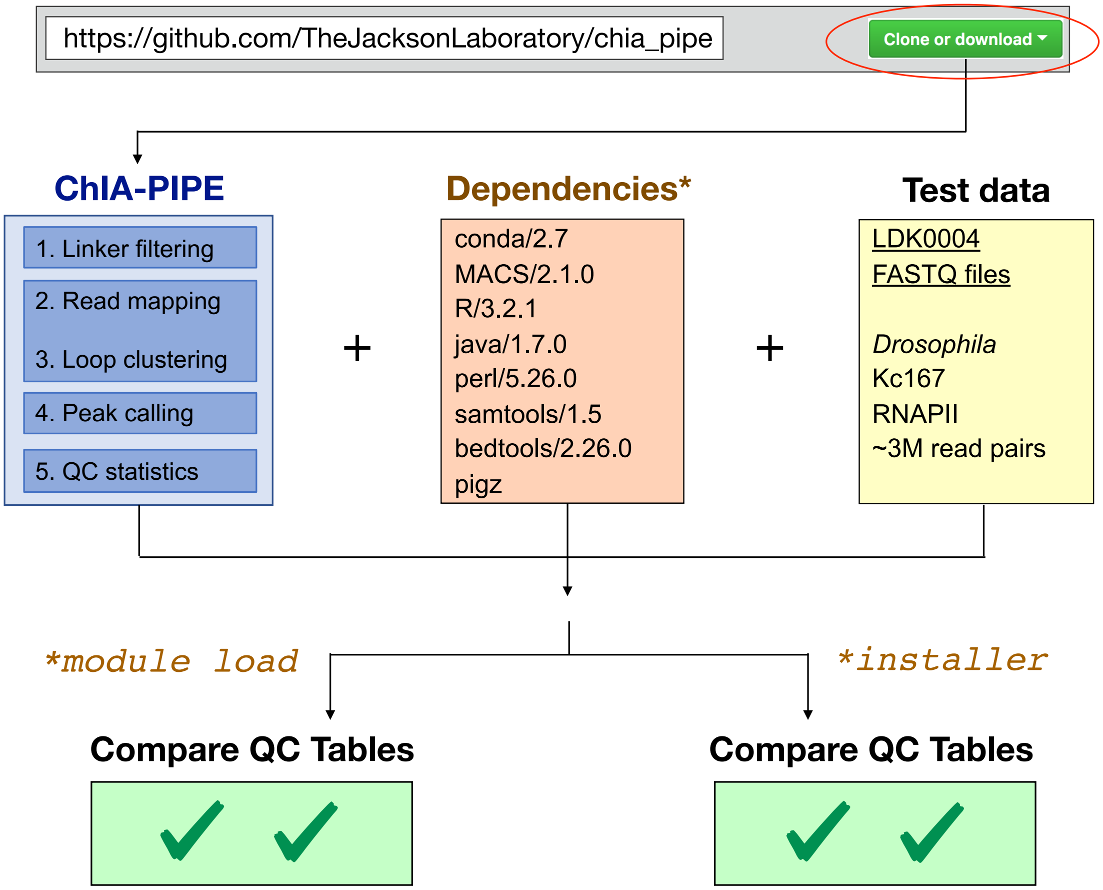


**Figure S1**. ChIA PIPE installation.

**Figure S1**. **ChIA PIPE installation**.

The ChIA PIPE source code can be downloaded from its Github repository. The dependencies of ChIA PIPE are listed in the orange box in the middle. In some high-performance computing clusters, the dependences can simply be loaded using the ‘module load’ command. Otherwise, ChIA-PIPE also includes an installation script for performing a local install of the dependencies. Finally, a small test set is provided of RNAPII ChIA-PET in *Drosophila* Kc167 cells. When this data set is processed, the resulting quality-control table can be compared to a reference quality-control table to ensure proper installation of the pipeline.

A

### ChIA-PIPE Quality-Control Table

| Library ID | LHH0054H | LHH0058H |
|---|---|---|
| Reference genome | hg38 | hg38 |
| Cell type | HFFc6 | HFFc6 |
| Factor | CTCF | RNAPII |
| Total Read Pairs | 345,288,614 | 360,358,387 |
| **Fraction read pairs with linker** | ***0.95*** | ***0.92*** |
| Read pairs with linker | 328,522,758 | 332,623,239 |
| **Fraction of linker reads with paired-end tags (PETs)** | ***0.60*** | ***0.62*** |
| PETs | 197,805,111 | 206,620,229 |
| **Fraction of PETs uniquely mapped** | ***0.72*** | ***0.76*** |
| Uniquely mapped PETs | 142,247,228 | 156,337,307 |
| Non-redundant PETs | 76,325,172 | 45,694,270 |
| **Peaks** | **23,413** | **35,518** |
| Intra-chrom PETs | 46,473,874 | 16,790,412 |
| **Ratio intra-to-inter -chrom PETs** | ***3.68*** | ***2.22*** |
| **Loops (PET count ≥ 3 and peak support)** | **34,162** | **45,214** |
| **Ratio of loops to inter-chrom PET clusters** | ***9.46*** | **2.64** |

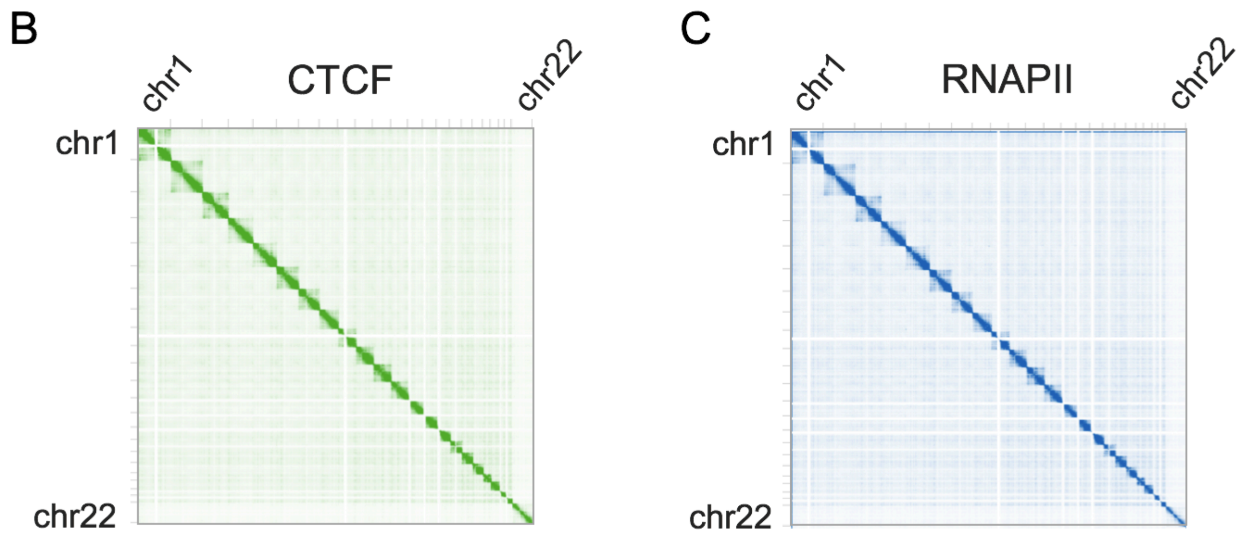


**Figure S2.** ChIA PIPE quality-control visualization.

**Figure S2. ChIA PIPE quality assessment and visualization.**

(A) An example quality assessment (QA) table generated by ChIA-PIPE. QA results are shown for one CTCF library and one RNAPII library in HFFc6 cells. Key parameters are: the fraction of read pairs with linkers (indicating ligation efficiency), the fraction of paired-end tags (indicating tagmentation optimization), the fraction of PETs uniquely mapped (indicating the amount of usable data), the number of called binding peaks, the number of loops, and the ratio of intra-to-inter chromosomal PETs and the ratio of intra-to-inter chromosomal loops (both indicating the signal to noise ratio of the data). (B-C) Full-genome Juicebox heat maps are shown for CTCF ChIA-PET (B) and RNAPII (C) ChIA-PET in HFFc6 cells (indicating the signal to noise ratio of the data).

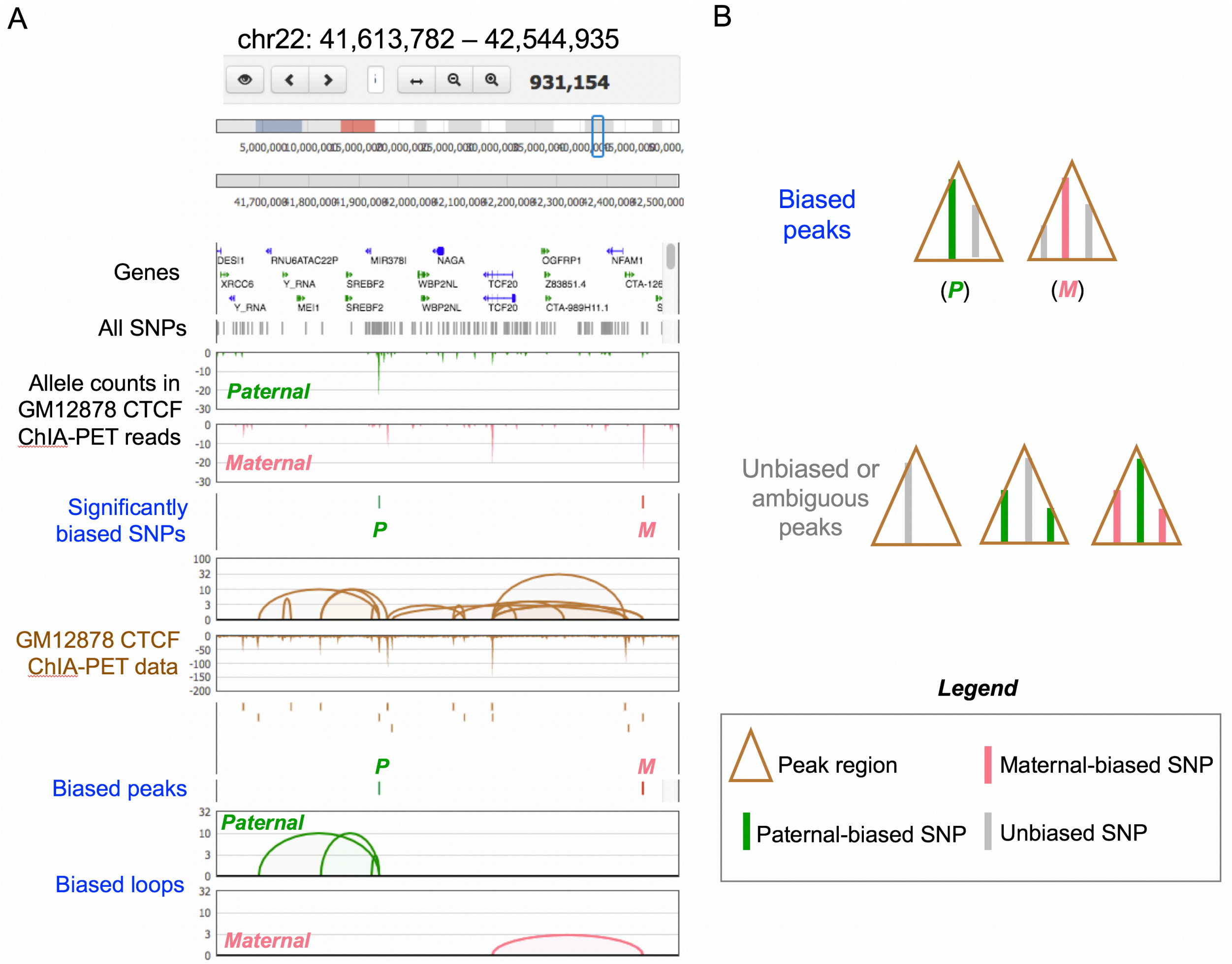


**Figure S3.** ChIA-PIPE resolves allele-specific chromatin interactions.

**Figure S3. ChIA-PIPE resolves allele-specific chromatin interactions.**

(A) An additional BASIC Browser example of a genomic region with one paternally biased SNP and one maternally biased SNP, and the corresponding biased ChIA-PET peaks and loops. (B) A diagram demonstrating how ChIA-PIPE categorizes peaks that overlap more than one SNP. In such cases, a peak is considered biased only if the SNPs are biased in the same direction (paternally biased SNPs and unbiased SNPs; or maternally biased SNPs and unbiased SNPs) and if the SNP with the highest read coverage in the peak is a biased SNP.

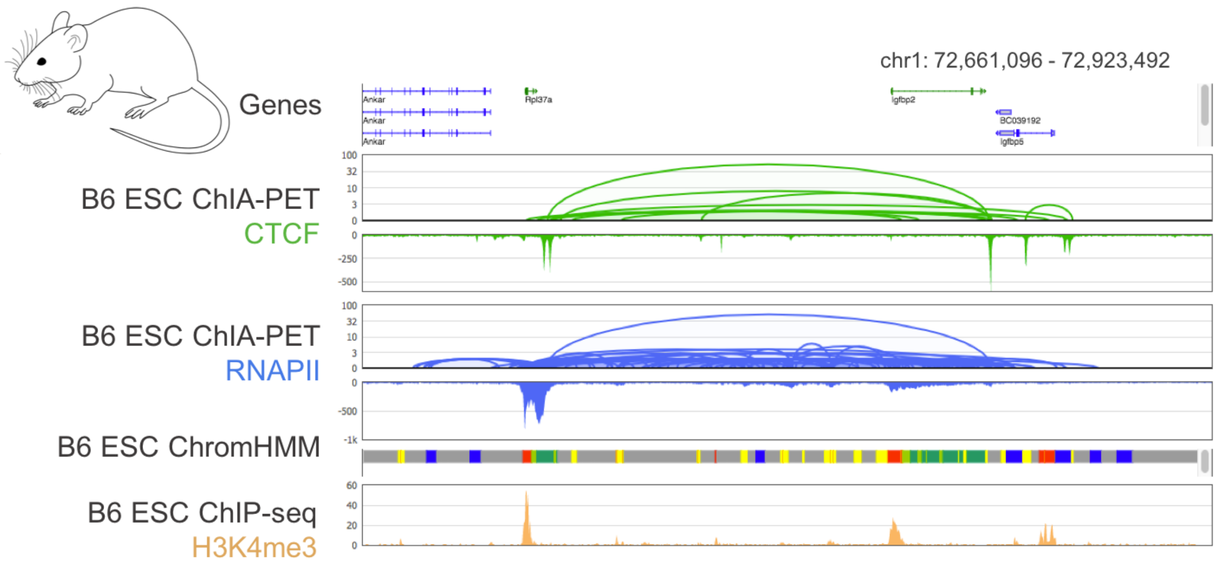


**Figure S4.** BASIC Browser for interactive, high-resolution ChIA-PET data visualization.

**Figure S4. BASIC Browser for interactive, high-resolution visualization of ChIA-PET loops, domains, and coverage.**

An additional BASIC Browser example of CTCF and RNAPII ChIA-PET data in mouse embryonic stem cells. Shown are the ChIA-PET loops and binding coverage. BASIC Browser supports the visualization of other data tracks to facilitate ChIA-PET interpretation. For example, the UCSC Known Genes are shown at the top and ChromHMM in the same cell type (from the ENCODE portal) and H3K4me3 ChIP-seq in the same cell type (from the 4DN) portal are shown at the bottom.

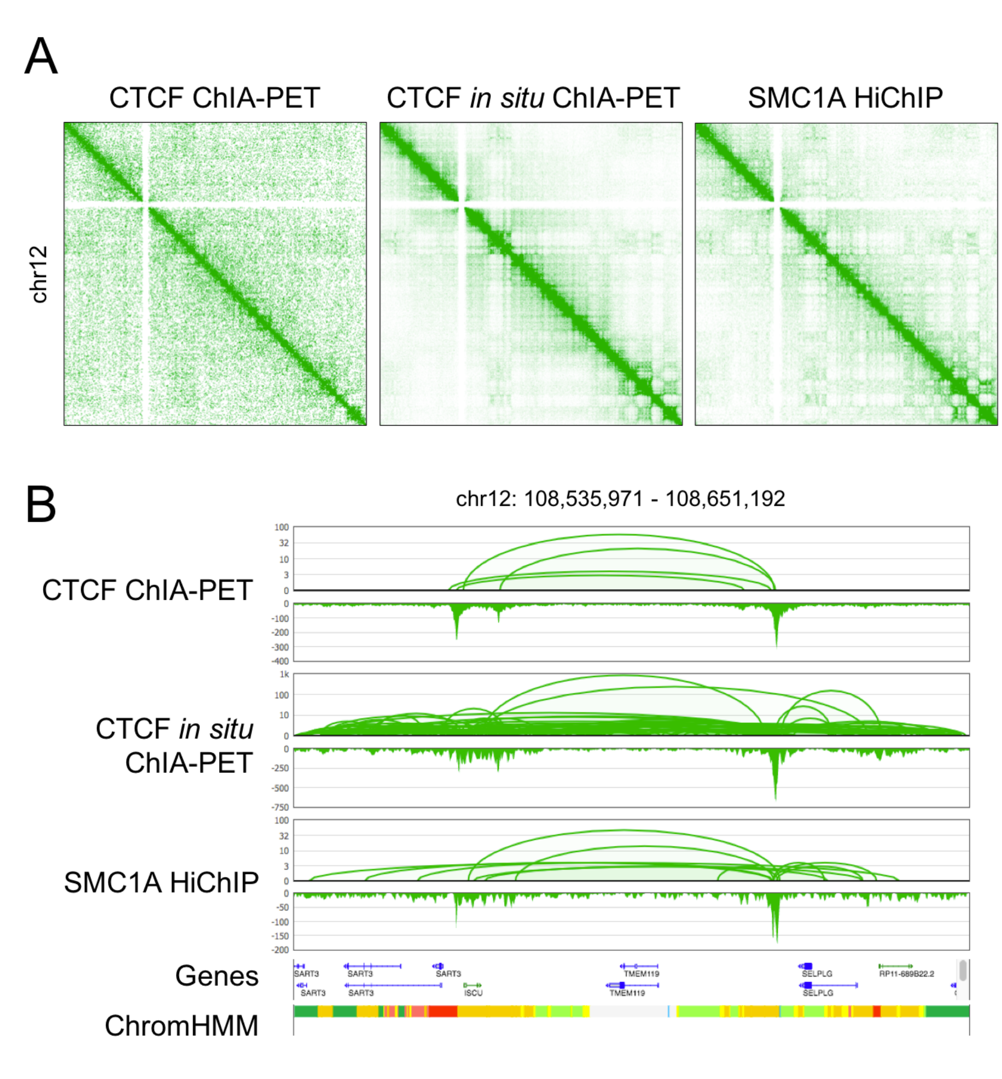


**Figure S5.** ChIA-PIPE can be used to process data from related 3D-genome mapping methods, such as HiChIP.

**Figure S5. ChIA-PIPE can be used to process data from related 3D-genome mapping methods, such as HiChIP.**

ChIA-PIPE was used to process comparable data sets in GM12878 cells from different methods: CTCF ChIA-PET, CTCF *in situ* ChIA-PET, and SMC1A (cohesin) HiChIP. ChIA-PIPE was readily adapted to process HiChIP data by treating the ligated restriction enzyme site as a “pseudo-linker”. (A) 2D heat maps of the three data sets from Juicebox.js. (B) BASIC Browser views of loops and binding coverage from the three data sets, alongside gene annotations and ChromHMM in GM12878 (from the ENCODE portal).
